# Monitoring lineages of growing and dividing bacteria reveals an inducible memory of *mar* operon expression

**DOI:** 10.1101/2022.09.16.508303

**Authors:** CC Guet, L Bruneaux, P Oikonomou, M Aldana, P Cluzel

## Abstract

In Gram negative bacteria, the *m*ultiple *a*ntibiotic *r*esistance or *mar* operon, is known to control the expression of multi-drug efflux genes that protect bacteria from a wide range of drugs. Since different drugs induce this response, identifying the parameters that govern the dynamics of its induction is crucial to better characterize the process of tolerance and resistance. Most experiments have assumed that the properties of the *mar* transcriptional network can be inferred from population measurements. However, measurements from an asynchronous population of cells can mask underlying phenotypic variations of single cells. We monitored the activity of the *mar* promoter in single *Escherichia coli* cells in linear micro-colonies and established that the response to a steady level of inducer was heterogeneous within individual colonies. Specifically, sub-lineages defined by contiguous daughter-cells exhibited similar promoter activity, whereas activity was greatly variable between different sub-lineages. Specific sub-trees of uniform promoter activity persisted over several generations. Statistical analyses of the lineages suggest that the presence of these sub-trees is the signature of an inducible memory of the promoter state that is transmitted from mother to daughter cells. This single-cell study reveals that the degree of epigenetic inheritance changes as a function of inducer concentration, suggesting that phenotypic inheritance may be an inducible phenotype.

## Introduction

Advances in time-lapse microscopy (Locke & Elowitz 2009) and substrate-printing (Bennett & Hasty 2009), combined with new fluorescent reporter proteins, have facilitated the characterization of stochastic phenotypic behaviors (Le et al 2005, Cluzel et al 2000, Elowitz & Leibler 2000, Guet et al 2008, Chang et al 2022) that are often masked by population measurements (Balaban et al 2004). Most of these methods depend upon measuring exclusively the concentration of fluorescent reporters. The accumulation and dilution of these reporters obscure time-dependent fluctuations at short timescales. These fluctuations are less of a limitation at long timescales in studies of bistable systems such as the *ara* and *lac* operons (Novick & Weiner 1957, Vilar et al 2003, Siegele and Hu 1997) or bacterial persistence (Balaban et al 2004), in which positive feedback loops lead to clearly distinct “on” and “off” states (i.e. digital networks). However, most gene regulatory networks produce a continuously varying distribution of outputs (i.e. analogue networks), and thus detecting the transmission of information from cell to cell especially at cell division can be very difficult (Taniguchi et al 2010, Austin et al 2006). In order to detect inheritance in networks with analog outputs, we have developed a technique using promoter activity time-series in single-cell lineages to quantify epigenetic inheritance for a gene regulatory network (Figure 1). We use the promoter of the *m*ulti*-a*ntibiotic *r*esistance (*mar*) operon from *Escherichia coli* fused to a gene coding for a fluorescent protein to test our technique. The study of single cell lineages reveals the presence of an inducible inheritance phenotype in the expression of the *mar* promoter as measured from a very low-copy reporter plasmid.

**Figure 1.**
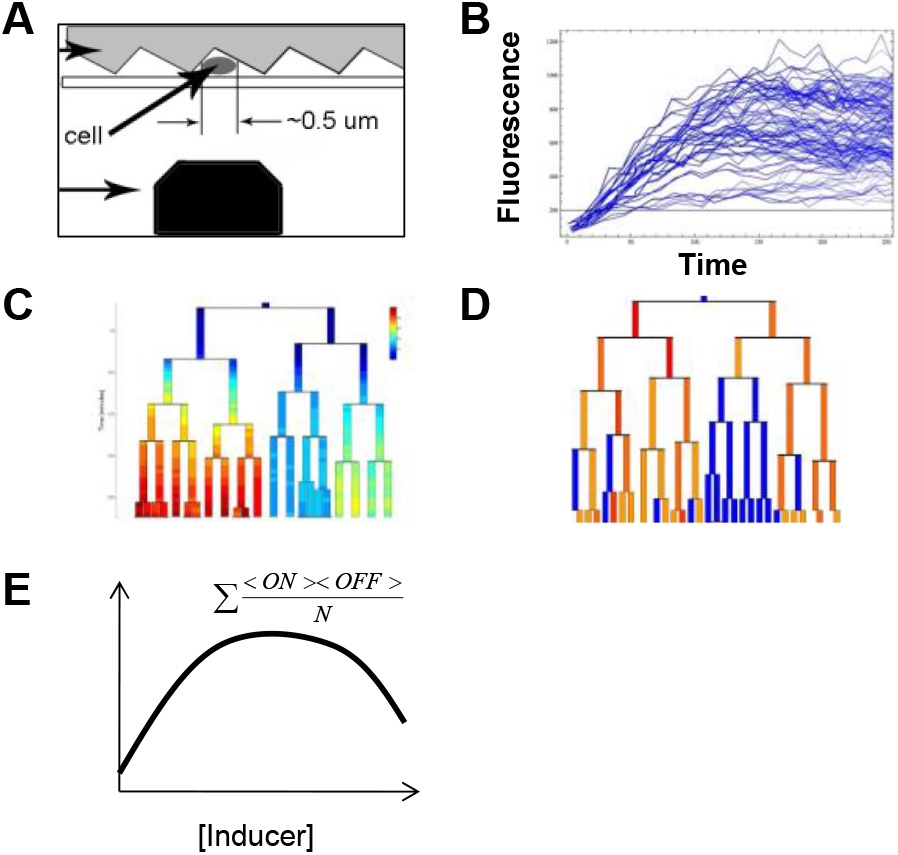
Schematic map of our experimental and analysis approach. **(A)** Data collection. Single *E*.*coli* cells are trapped between a patterned agarose pad and cover slip, and are excited via a blue laser and scanning nano-positioning stage. **(B)** Conversion into time series of total fluorescence. From the raw data, time series of fluroescence are produced across growing colonies. **(C)** Calculation of promoter activity across individual cell cycles. These colonies are analyzed and broken up into individual cell-lineages, with the size and concentration of fluorescent reporter given at each time point for each cell, from which cells are then given a single value for promoter activity between segmentation events. **(D)** Binarization of promoter activities. Using the distribution of these promoter activity values, cells are then given a value of <ON> or <OFF>. **(E)** Calculation of domain size product. The domains of contiguous ON or OFF cells are measured, and multiplied to determine their overall measure of inheritance.

The *mar* operon also denoted as *marRAB*, consists of three genes: *marR* - a repressor, *marA* - an activator and *marB* - whose function is not fully understood. The biology of the *mar* operon has been extensively studied at the population level (Martin et al 1996), yet little is known about the dynamical aspects of this inducible gene regulatory network at the single-cell level. Through the direct and indirect action of MarA on about 60 different genes, *mar* represents one of the largest regulons of *E. coli* (Barbosa & Pomposiello 2005). One of the main targets of MarA is activation of the expression of the multi-drug resistance efflux pumps operon *acrAB* (Li & Nikaido 2004). In the absence of an inducer, the *mar* network is characterized by a negative feedback loop produced by MarR self-repressing the *mar* operon (Alekshun & Levy 1997) (Figure 2). In the presence of an inducer, such as salicylate, MarR repression is eliminated, and the activator MarA initiates its own expression, constituting a positive feedback loop. MarA also activates the *acrAB* and *tolC* operons, that together encode the AcrAB-TolC efflux pumps)Martin et al 2008), the main multi-drug resistance determinant of Gram negative bacteria. Over time, the accumulation of efflux pumps reduces the intracellular concentration of inducer, which constitutes a negative feedback loop by renewing the repression of the *mar* operon by MarR. Thus, the activation of the *mar* system confers multi-drug resistance (George & Levy 1983) that is eliminated when the pumps are no longer expressed (Alekshun & Levy 1997, Le et al 2006). The reversible nature of this resistance suggests that individual cells may have widely ranging responses to a steady input (Lewis 2007). Interestingly, the alternating behavior of the positive and negative feedback loops (Figure 2) has the potential to reverse the activation of the *mar* promoter even under a steady level of inducer (Le et al 2006, Thomas et al 1988, Aracena 2008).

**Figure 2.**
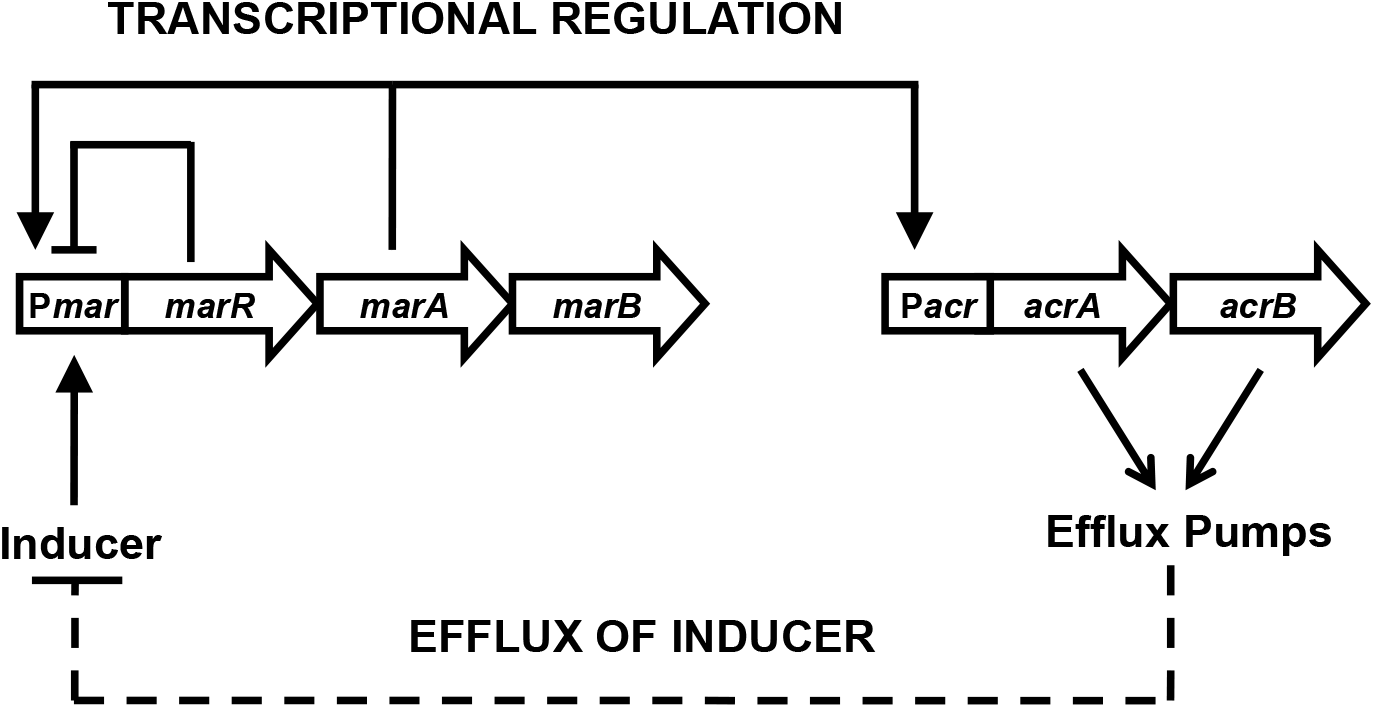
Topology of the networks formed by the *marRAB* and *acrAB* operons. The core transcriptional network of the *mar* system includes the repressor MarR that regulates the *marRAB* operon. In the presence of the inducer salicylate, MarR repression is abolished, allowing MarA to activate its own expression and the expression of the *acrAB* operon. The products of the *acrAB* genes form the efflux pump complex AcrAB-TolC. Efflux pumps create negative feedback on *marRAB* activity due to efflux of the inducer salicylate.

Here, we measured activity of the *mar* promoter in single cells using a confocal scanning volume across *E. coli* colonies growing linearly, starting from a single ‘mother’ cell. The *mar* promoter controls the expression of a *yfp-venus* reporter with very fast maturation time (Balleza et al 2018) from a very low-copy plasmid with SC101* *ori* (Lutz & Bujard 1997). Using the promoter activity reported via YFP expression, we constructed genealogy trees describing promoter induction for individual lineages of *E. coli*. We used an algorithmic process to binarize the output of the promoter and then calculated a measure for the degree of epigenetic inheritance of each individual lineage at various inducer concentrations (Figure 1). Surprisingly, we found that the coefficient of variation (CV) of expression depended in a non-monotonic way on inducer concentration. Based on our analysis, we found that the degree of epigenetic inheritance changed as a function of inducer concentration, suggesting that phenotypic inheritance may be an inducible phenotype within an analog-output gene regulatory network.

## Materials and Methods

### Strains and plasmids

We used the *E. coli* K-12 strain Frag1B (Le et al 2005) and JW5503-1, a *ΔtolC* deletion mutant (Baba et al 2006). Plasmid *pZS*1mar-venus*: We replaced the *gfp* from pZS*1*R*-*gfp* (Guet et al 2002) with the *venus-yfp* gene using KpnI and HindIII sites. We PCR amplified the *mar* promoter, containing the marbox and the two MarR operators, from the Frag1B chromosome. Since one operator for MarR overlaps with the *marR* coding region, we retained the first seven amino acids at the N-terminus of MarR and fused them to Venus-YFP (Martin et al 2004). We mutated the wild-type GTG start codon of *marR* to ATG by changing G with A in the primer. We cloned the *mar* promoter into the XhoI and KpnI sites of pZS*1*R*-*venus*. The KpnI site introduced a Gly-Thr linker between MarR and Venus-YFP. The pZS*1*mar-venus* plasmid contains Amp^R^ and has a SC101* *ori* that maintains the copy number at 3-4 copies/cell (Lutz & Bujard 1997). We transformed all strains by electroporation.

### Growth conditions

We grew cells overnight at 30°C in Luria-Bertani broth (LB) containing 100μg/ml ampicillin (Sigma Aldrich, St. Louis, MO, USA). We diluted the overnight cultures 1:600 in fresh LB media and harvested cells after an additional 3.5 hours at 30°C. The optical density (OD) of the final cultures varied between 0.12-0.15 as measured at 600nm.

### Sample preparation for microscopy and submicron-groove fabrication

We prepared fresh 3% low melting agarose (Fisher Scientific, Pittsburgh, PA, USA) in LB in a 70°C water bath. We mixed salicylate from a 1M sodium salicylate (Fluka, Switzerland) stock solution with 100 μL LB and added the mixture to 5 mL LB-agarose and swirled vigorously for 1 min. in order to achieve rapid and homogeneous mixing. We poured 100 μL of this 3% agarose-LB as a 1 cm^2^ gel slab onto an optical grating (300 lines/mm, blaze angle 26°, Bausch and Lomb, Rochester, NY, USA). The gel pad solidified for 20 min in the dark at room temperature, imprinting the grooves of the optical grating in the gel. We placed 0.3 μL of freshly harvested *E. coli* cells on top of a thin coverslip (Fisher Scientific, USA) sealed with wax to an aluminum holder. We placed the agarose-LB slab on top of the 0.3 μL droplet of cells. Due to surface tension, most cells aligned themselves in the submicron-grooves. The sample was sealed and immediately placed on the heating stage of an inverted Olympus X71 microscope (Olympus, Japan). We began collecting data within 5-10 min of the sample set up.

### Data collection and processing

To measure fluorescence from individual cells, we focused blue laser light (Sapphire 488 nm, 20 mW, Coherent, Santa Clara, CA, USA) into a diffraction-limited spot (width = 0.4 μm) (Guet et al 2008) and scanned cells at 4 μm/sec using a nano-positioning stage (Physik Instrumente, Auburn, MA, USA). Emitted photons were collected in a confocal geometry and detected with an avalanche photodiode (ALV, Langen, Germany). This scanning procedure is minimally invasive and accurately measures fluorescence intensity within individual cells (Guet et al 2008). We scanned each cell every 5-10 min. We aligned the scanning trajectory of the laser spot perfectly along the submicron-grooves using the nano-positioning stage. Fluorescence data was stored for further analysis.

Each scan produced a fluorescence profile of individual cells from a micro-colony. Fluorescent data was binned into 0.4 μm pixels. Because the excitation spot was small (width = 0.4 μm), cell cleavage sites appeared as sharp dips in the fluorescence profile while cell bodies appeared as long, bright fluorescence plateaus (see Figure 1B). Cell length was the distance between two consecutive cleavage sites. The difference in length between two consecutive measurements was used to estimate the cell growth rate. To produce genealogy trees, we used a visual interface (in house software, Matlab). Using this program, we manually picked cleavage sites from the intensity profiles produced by the scanning and assigned and tracked cell lineages. The intracellular concentration of fluorescent proteins was given by the brightest pixel. The fluorescence intensity was directly related to the intracellular concentration of fluorescent molecules. The promoter activity was the change in fluorescence intensity between two time points corrected for by a dilution factor that accounts for growth. We assigned each cell a single value for promoter activity between divisions.

### Data analysis

*Binarization:* We binarized promoter activity as either ON (active) or OFF (inactive) using a threshold defined in Figure 3A. Activity was ON if the cell had promoter activity at least two standard deviations above the mean of the distribution of activities of un-induced cells. Activity was OFF if the promoter activity was below this threshold.

**Figure 3.**
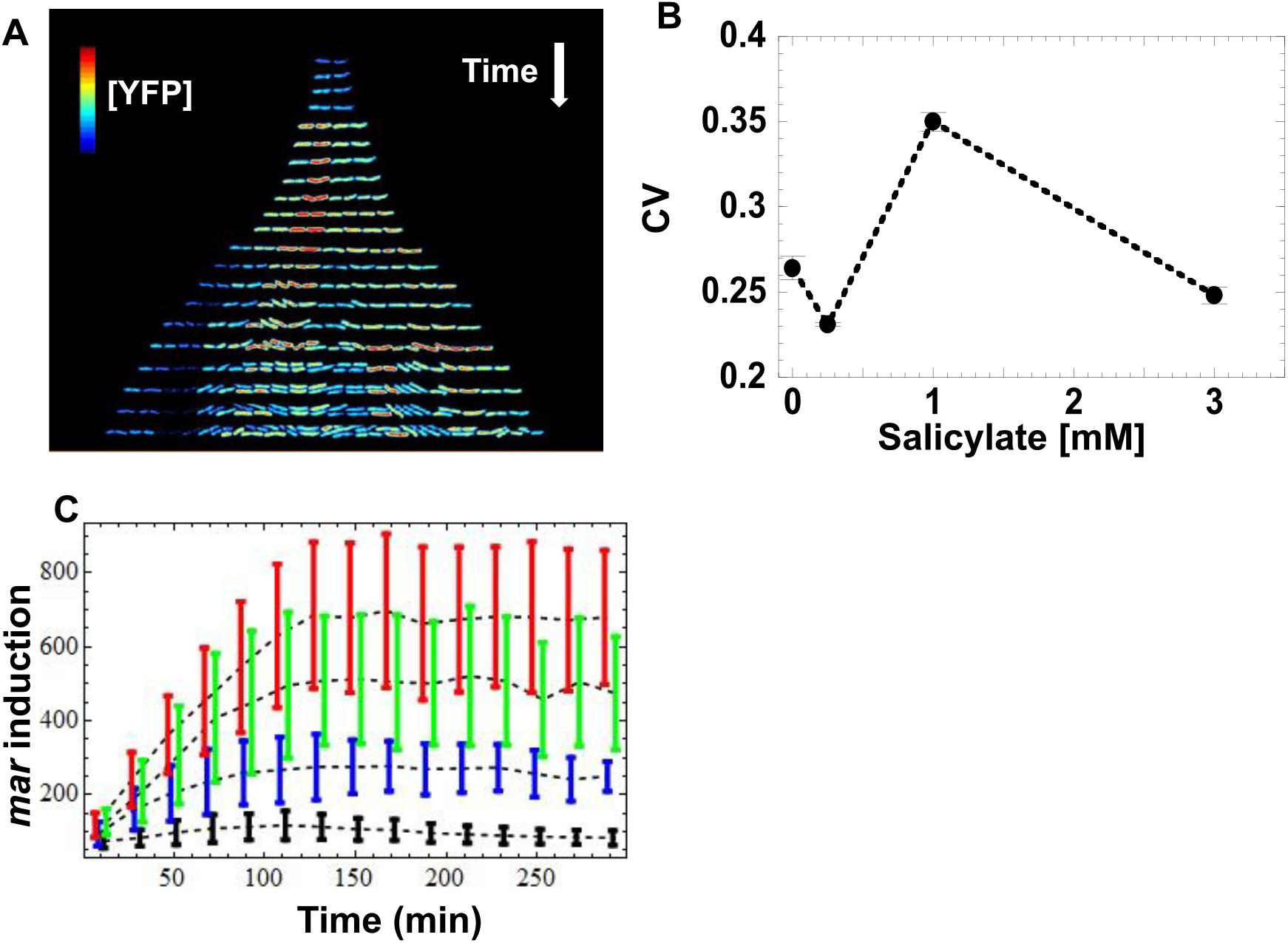
Induction of *mar* promoter with salicylate. **(A)** Induction of the *mar* promoter in individual cells of a linear colony. Single colony images are separated by ∼20 minutes. The heat map shows fluorescence intensity (Red = High, Blue = Low) from YFP expression driven by the *mar* promoter activity in individual cells exposed to a steady level of salicylate (1 mM). Induction starts at t = 9 min. **(B)** Mean coefficient of variation of [YFP]. YFP fluorescence from single cells was averaged for 10-minute intervals and the coefficient of variation across the interval was calculated. The mean coefficient of variation across the interval 120 to 300 minutes was then calculated. Error bars represent standard error. **(C)** Induction kinetics of the *mar* promoter at the population level. YFP fluorescence from single cells was averaged for 10-minute intervals, reporting *mar* promoter activity as a function of time. Four different salicylate concentrations were used: 0 mM (black), 0.25 mM (blue), 1 mM (green), 3 mM (red). Error-bars are the standard deviation from the distribution of single cell measurements.

### Degree of Inheritance

For each tree, we defined a sub-tree as two or more cells that were directly connected (through mother-daughter or sister-sister relationships) and all shared the same promoter activity, i.e. ON or OFF. For each tree, we computed the mean sub-tree size of ON and OFF sub-trees, *<ON>* and *<OFF>*, respectively. The mean sub-tree size is a measure of how large the average sub-tree is in which a randomly selected cell finds itself on a genealogy tree. It is defined as:

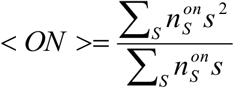, where 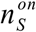 is the number of ON domains that contain *s* cells, and represents the definition of a mean domain size in percolation theory (Stauffer & Aharony 1994). For each tree we calculated the product *<ON><OFF>*. For a given inducer concentration, the degree of inheritance was defined as the average domain size product, or ∑< *ON* >< *OFF* > divided by the number of trees, *N*.

### Computational model

We implemented a discrete-time branching process in which all cells in the population divide simultaneously. Each cell can be in one of two possible states: active (on) or inactive (off). When a cell divides, each of its two daughter cells can be active or inactive depending on the state of the mother cell. To implement the model, four conditional probabilities must be defined. Denoting as DC and MC the daughter and mother cells, respectively, the four conditional probabilities are: *P*(*DC on*| *MC on*), *P*(*DC off*|*MC on*), *P*(*DCon*|*MC off*), and *P*(*DC off*|*MC off*). Only two of these four probabilities are independent since, *P*(*DC on*|*MC on*) = 1 − *P*(*DC off*|*MC on*), and *P*(*DC off*|*MC off*) = 1 − *P*(*DC on*|*MC off*). Therefore, the complete branching process is thoroughly characterized by the two probabilities *P*(*DC off*|*MC on*) and *P*(*DCon*|*MC off*). These two probabilities were computed from experimental data for different inducer concentrations. Once the relevant probabilities are fixed, the branching process starts with one cell in the “on” state assuming that this cell has been placed in a medium with nonzero inducer concentration. From there, the branching process evolves.

## Results

### Constructing genealogy trees of promoter induction in wild-type cells

We cloned the full length *mar* promoter upstream of Venus *yfp* (Nagai et al 2002), on a very low copy plasmid (3-4 copies per cell (Lutz & Bujard 1997). The fast maturation of the Venus-YFP chromophore allows us to report (Balleza et al 2018) changes in *mar* promoter activity within a few minutes. We monitored salicylate-induced expression of the *mar* promoter as a function of time in wild-type *E. coli* cells at the single cell-level across several inducer concentrations that cover the dynamic range of previous population measurements (Rosner 1985, Cohen et al 1993). To limit cell-cell contact, we restricted cell growth to one dimension by growing bacteria as linear colonies within long micro-grooves imprinted into agar. This geometry ensures that all cells are grown in similar micro-environmental conditions. To ensure the genetic homogeneity of a linear colony during growth, we restricted our measurements to micro-grooves that were initially occupied by only a single bacterium. By scanning the colonies with a focused blue laser and collecting the emitted fluorescence with a photodiode (Guet et al 2008), we monitored the expression level of YFP in single cells exposed to a steady level of the inducer salicylate ranging from 0 mM to 3 mM. We analyzed the distribution of the concentration of the YFP reporter, [YFP], at 5-10 minute intervals in single cells across colonies. At the single-cell level, we observed that [YFP] varied several-fold within colonies, even between siblings (Figure 3A). Using the coefficient of variation (CV) as a quantitative estimate of the cell-to-cell variability, we found the highest variability in [YFP] at 1 mM salicylate (CV = 0.35 at 1 mM; CV ≤ 0.25 for the other inducer concentrations) (Figure 3B). By contrast, the average [YFP] across colonies reached a well-defined steady state for each concentration of inducer about 100 min after induction (Figure 3C).

Next, we set out to determine whether this variation was a result of random fluctuations, or due to a transmission of cytoplasmic information from a mother cell to its daughters. To characterize how *mar* promoter activity changed through cell lineages, we monitored [YFP] across generations within the linear colonies and we represent the changes in activity using genealogy trees (Figure 4). We generated the trees by presenting the time series of [YFP] concentration for a single lineage under each inducer level, with the concentration of [YFP] denoted from low (blue) to high (red). Both, the absence of salicylate (0 mM) and high (3 mM) concentration of salicylate led to homogeneous [YFP] across generations (Figure 4, Fig S1b-h). Remarkably, an intermediate salicylate concentration (1mM) resulted in heterogeneous distribution of [YFP], ranging from very high (as observed with 3 mM salicylate) to very low (as observed with no salicylate) (Figure 4, Fig. S1f).

**Figure 4.**
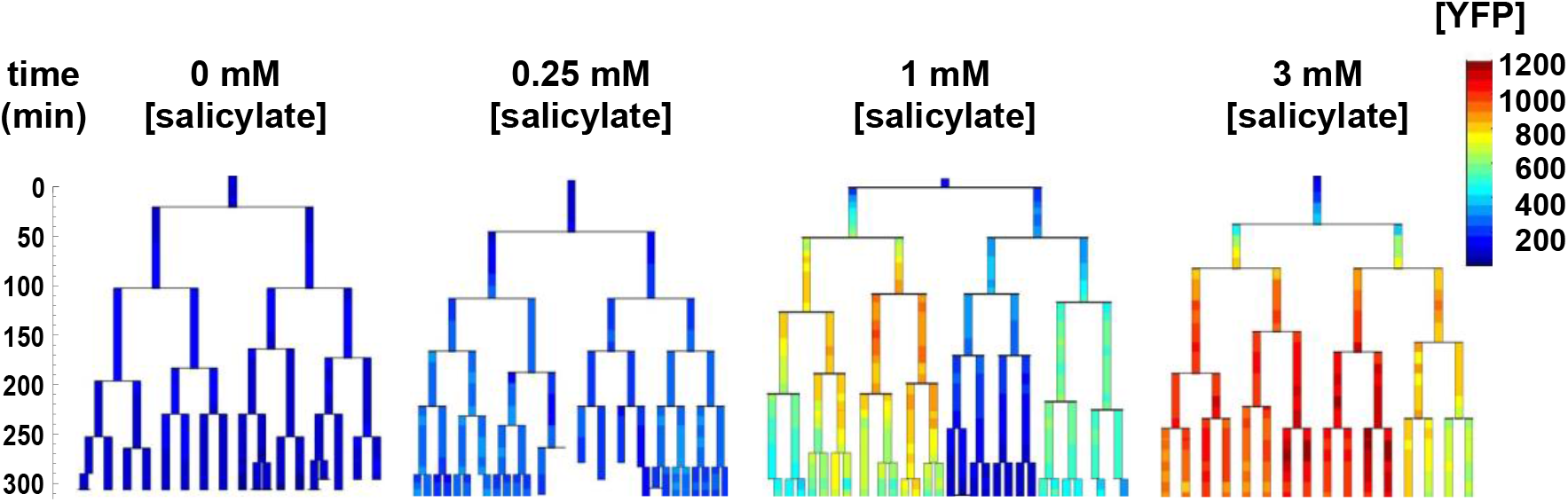
Genealogy trees of *mar* promoter induction in wild-type cells. Single cells with a plasmid carrying *yfp-venus* regulated by the *mar* promoter were monitored through several generations in the presence of the inducer salicylate. The genealogy tree follows the induced level of YFP concentration through the generations of a colony. Constant salicylate exposure began at time *t* = 0. Heat-map scale uses arbitrary fluorescence units to show low (blue) to high (red) [YFP].

### Binarization of promoter activity and quantification of inheritance

We estimated *mar* promoter activity by monitoring temporal variations of [YFP] while accounting for changes in cell volume due to growth (Figure 3A). To simplify our analysis, we used a threshold to binarize *mar* promoter activities as active or inactive within genealogy trees. In the non-induced condition, the *mar* promoter exhibited a natural leakiness (characteristic of most bacterial promoters) that produced a basal level of promoter activity. We chose a threshold that discriminates the distribution of non-induced cells (0 mM salicylate) from highly induced cells (3 mM salicylate) (Figure 5A). Cells with promoter activities below the threshold had an inactive *mar* promoter, while those with activities above the threshold had an active *mar* promoter. This binarization allowed for simple counting of active cells and inactive cell sub-trees (Figure 5B). To quantify this heterogeneity, we calculated the product of the average sizes of active and inactive sub-trees in a genealogy tree. If the average size of either active or inactive sub-trees is close to zero, then the degree of inheritance is small. When both active and inactive sub-trees are large, the degree of inheritance is high. We found that observed inheritance was highest in genealogy trees with cells induced at intermediate levels (Figure 5C). This measure was insensitive to specific thresholds (Figure 6).

**Figure 5.**
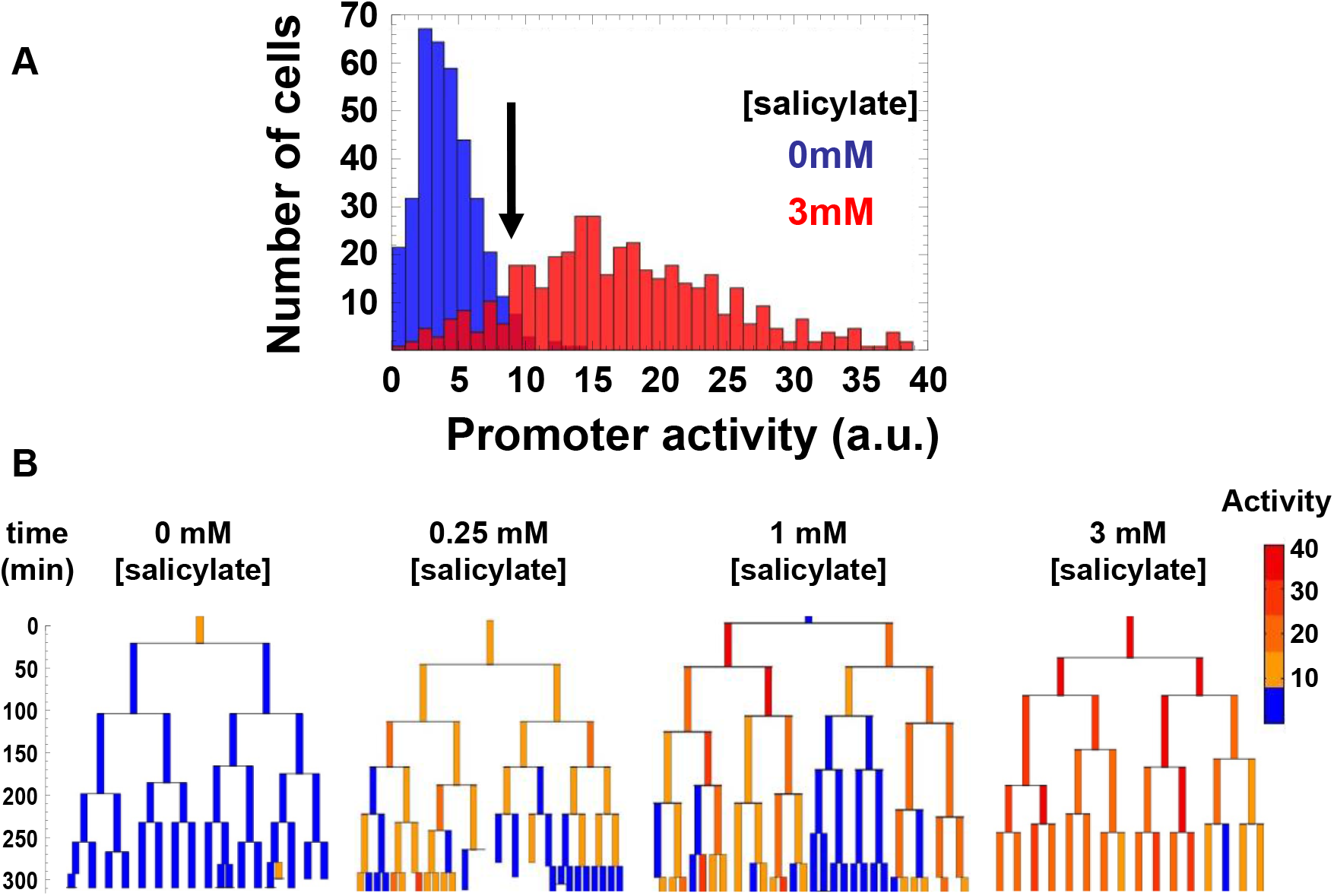
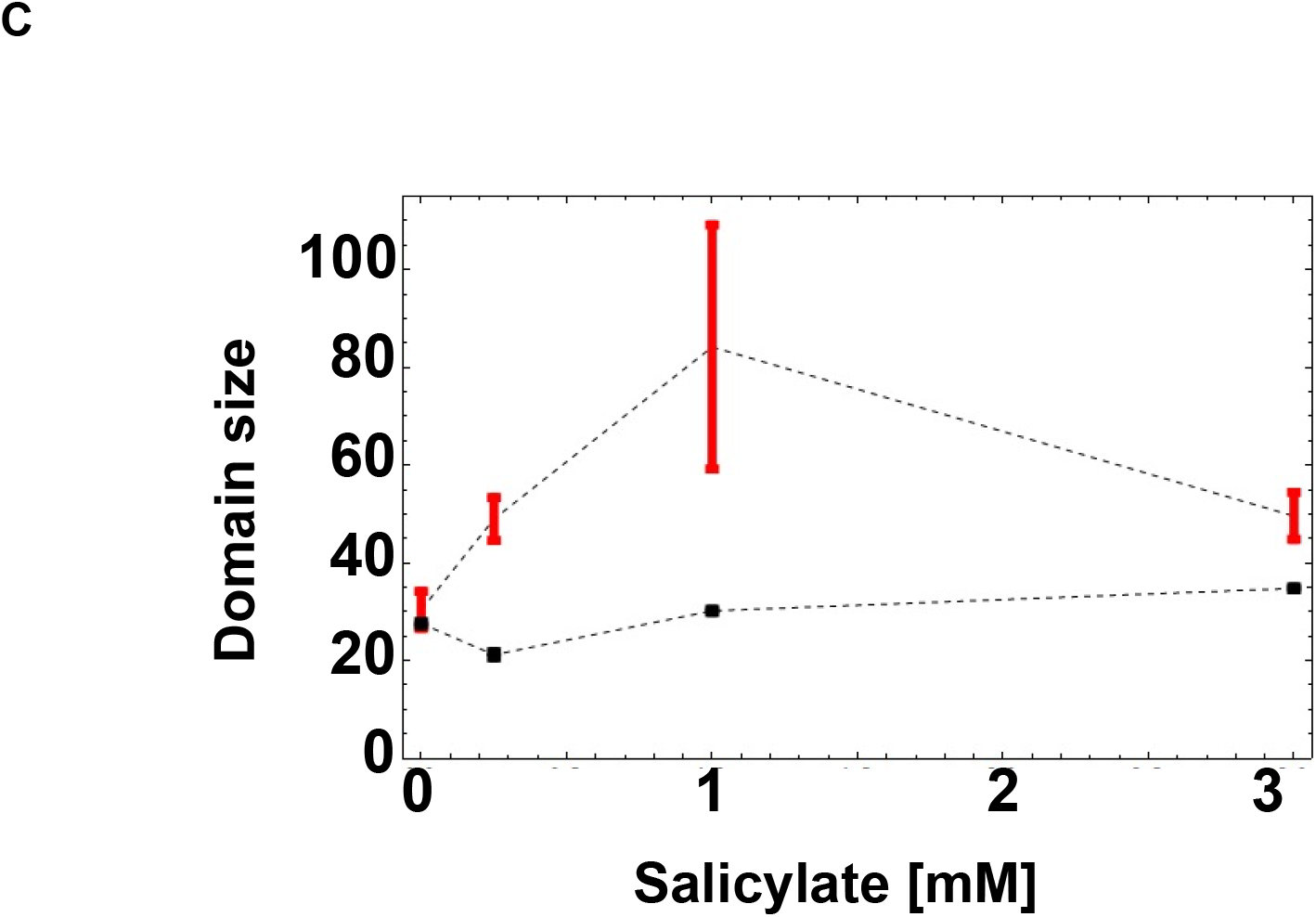
Binarization of the *mar* promoter activity. **(A)** Distribution of *mar* promoter activity across a population of non-induced 0 mM salicylate (blue) and fully induced 3 mM salicylate (red) cells. The promoter activity is estimated as the temporal derivative of the measured YFP concentration corrected for cell growth. The black arrow shows the threshold between active and inactive promoter states defined at two standard deviations from the mean of the non-induced cell distribution (blue). **(B)** Genealogy trees of *mar* promoter activity for different inducer concentrations. The vertical axis represents time in minutes. To binarize the promoter activity we used the threshold from (A). Below the threshold, the *mar* promoter is inactive (blue), above the threshold the promoter is active (yellow, orange, red). **(C)** Domain Size Product of trees is shown for wild-type genealogy trees (circles) and randomly reshuffled genealogy trees (squares) at different inducer concentrations. Error-bars are the standard error.

**Figure 6.**
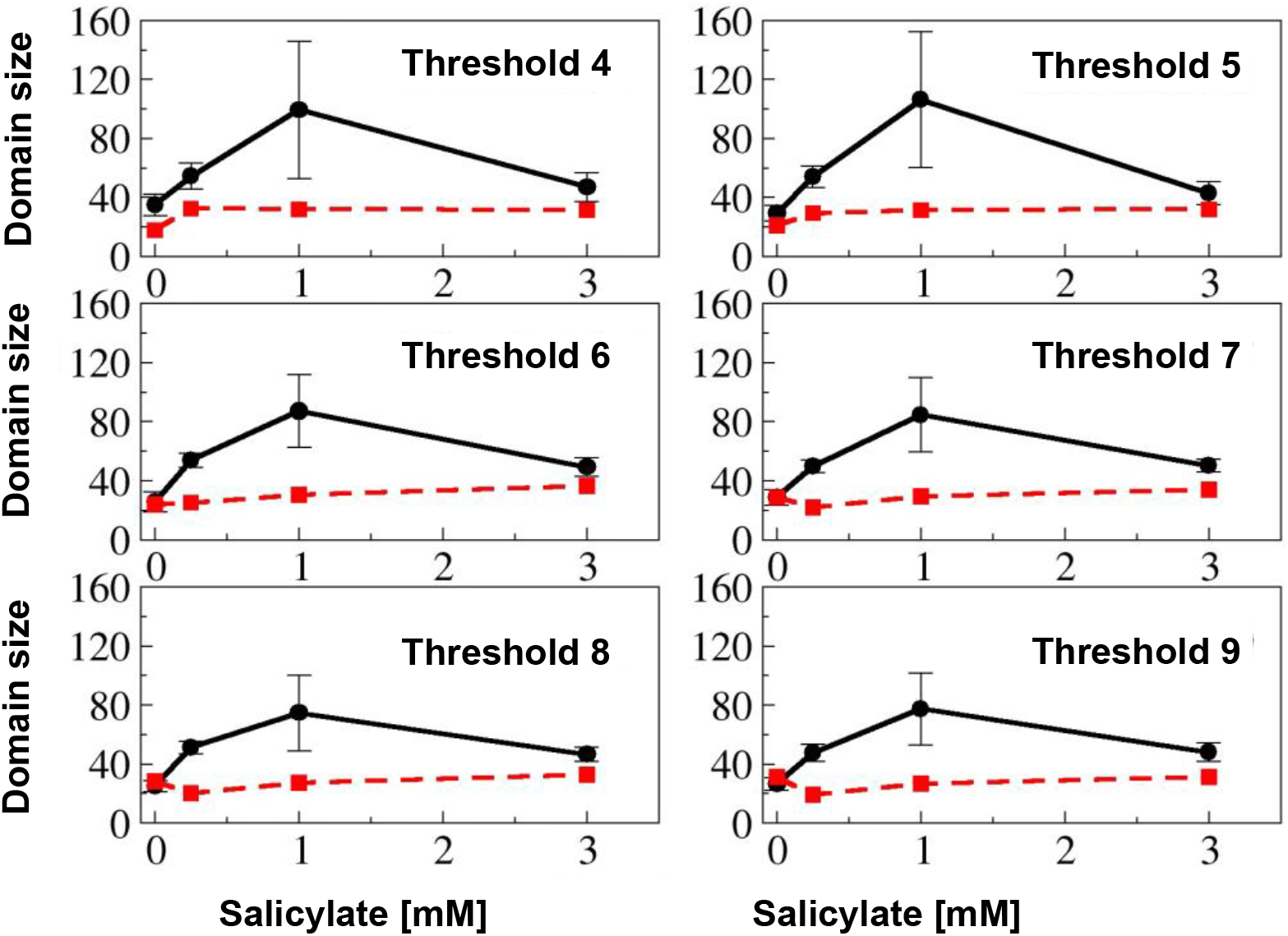
Epigenetic memory dependence on threshold choice. Plots of the branch size product as a function of the inducer concentration for different choices of the binarization threshold. The curves correspond to wild-type genealogy trees (black lines with circles) and randomly reshuffled genealogy trees (red lines dashed with squares). The error-bars are the standard error. Note that in all cases the maximum branch size product occurs at 1mM regardless of the binarization threshold used.

### Observed inheritance is not a result of random switching

If promoter activity switches randomly at each cell division, then the activity across trees would exhibit a noisy distribution. Alternatively, if cells inherit the ancestral promoter activity state, then trees would display large sub-trees of similar promoter activity. To test if random switching alone can explain the observed promoter activity distribution, we generated randomized activity trees and calculated their domain size product. In order to render the theoretical result of the genealogy trees as relevant as possible to the experimentally obtained trees, we constructed each tree by randomly placing active and inactive cells on the leaves of the tree while keeping the number of active and inactive cells identical to those from the empirical genealogy trees. For all non-zero inducer concentrations, the randomized trees had a similar domain size product (Figure 7) that was always lower than that of the experimental trees (Figure 5C). These results suggest that information is being passed from the mother cell to the daughter cells, indicating the presence of epigenetic inheritance. One of the main feedback loops in the gene regulatory system we study is represented by the AcrAB-TolC efflux pumps that extrude inducer and thus their activity represents a negative feedback loop on the induction of the *mar* operon (Figure 2). A mutant *ΔtolC* mutant exhibited homogeneous [YFP] across all lineages and all inducer levels used (Figure S2). Thus the presence of this negative feedback look that acts at a longer time scale is essential for the process of inheritance of *mar* promoter activity.

**Figure 7.**
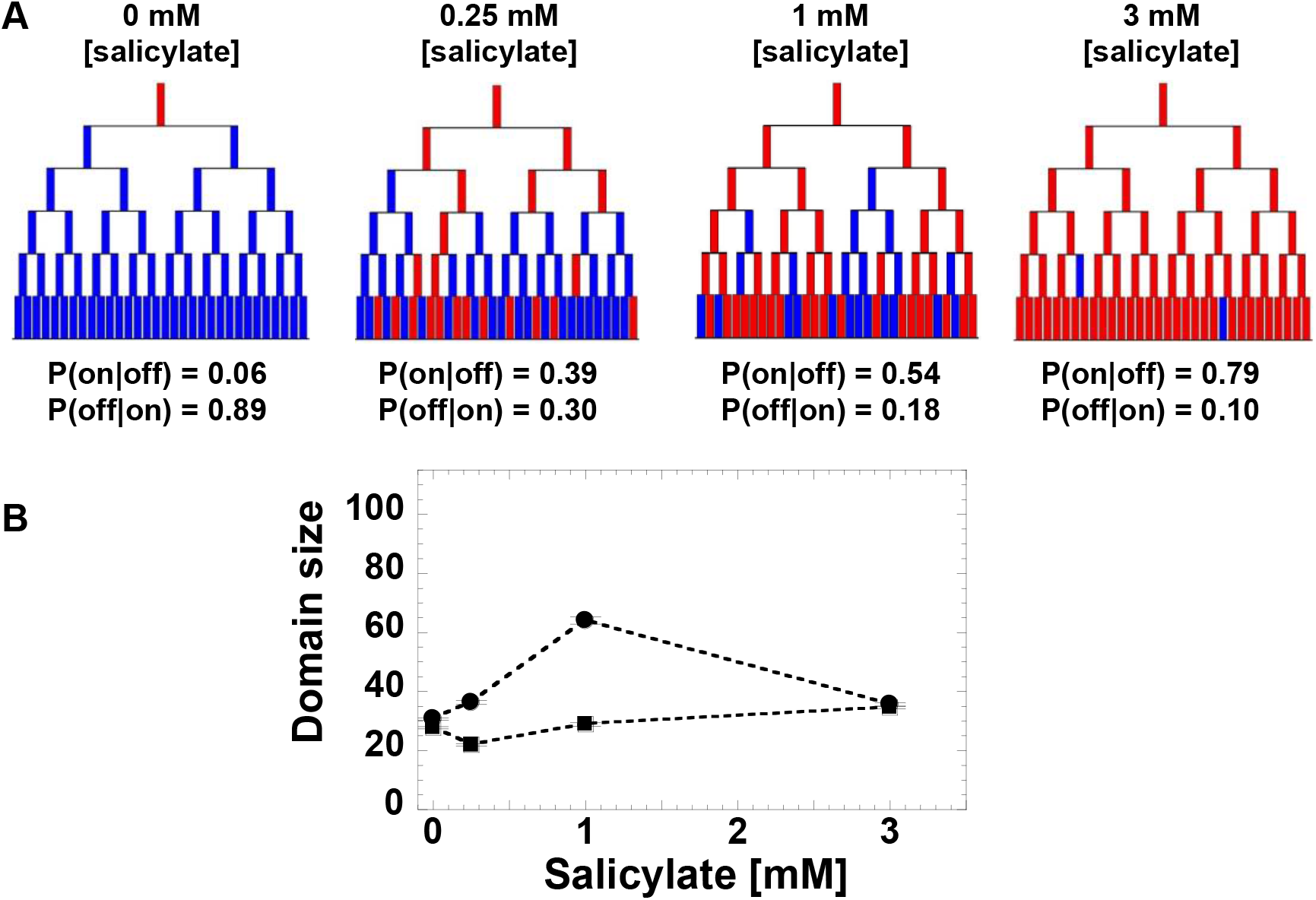
Toy model of the *mar* promoter activity. **(A)** Toy model of *mar* operon induction. Cells can switch between active and non-active states with conditional transition probabilities P(on|off) and P(off|on). Values of the probabilities were determined from the experimental data. The genealogy trees follow cells through generations and illustrate switching between an inactive (blue) and active (red) promoter. **(B)** Domain Size Product of trees was computed from 200 model simulations using the experimentally-derived conditional probabilities, P(on|off) and P(off|on) (circles), and non-conditional probabilities, P(on) and P(off) when the inheritance strength is zero (squares).

## Discussion

Our experimental technique has shown that phenotypic inheritance is also a measureable quantity in gene regulatory networks with analog output, and not only in bistable genetic networks. We have demonstrated that inheritance may be sensitive to conditions, such as the induction level in the *mar* operon, which remains undetectable when measurements are performed at the population level (Cohen et al 1993). The standard picture of the transcriptional regulation by MarR and MarA of the *marRAB* and *acrAB* operons has been established mostly through population level experiments (Martin et al 1996). Based on these experiments and our own population-level measurements, we hypothesized that, under steady level of induction, the *mar* promoter from single cells would also exhibit a steady level of activity. Surprisingly, our experimental approach showed that even well after the promoter reaches its steady-state value measured at the population level (∼100 min post induction), some single cells had an inactive *mar* promoter. Monitoring promoter activity through multiple generations revealed an unusual pattern of activity in induced cells within each isolated bacterial colony. We found that within lineages, cells with similar promoter activity form sub-trees across generations, suggesting an inducible inheritance where the activity of daughter cells reflects the promoter activity of the mother cell. Surprisingly, the level of inheritance peaks for an intermediate inducer concentration of 1mM in *E*.*coli* cells.

This inducible inheritance is more subtle and transient than in all-or-none examples, such as the *ara* and *lac* operons and bacterial persistence and has therefore been more elusive and difficult to trace. Here, by measuring promoter activity across lineages and by simplifying the output by means of binarization, we were able to measure the degree of cell-to-cell inheritance. This technique should be broadly applicable, to a number of gene regulatory networks, using readily available plasmid reporter libraries (Zaslaver et al 2006), e.g. flagellar assembly cascade, ribosomal assembly and metabolic networks. We anticipate that this single cell approach could be applied at a high-throughput level using promoter libraries, wide-field fluorescence microscopy, and automated image-processing software. Using this type of systems approach could reveal heretofore unknown inheritance phenotypes in the gene regulatory networks of *E. coli* and other single-celled organisms.

What can be the mechanistic origin of the inheritance we uncovered through our experimental and theoretical analysis? For one, the complex network of negative and positive feedbacks that the *marRAB* and *acrAB* operons form (Figure 2) is by its very topological construction prone to complex dynamics behavior. What adds extra complexity to this already complex dynamics, is the recently discovered biased partitioning of the AcrAB-TolC complex at old poles (Bergmiller et al 2017), which in itself is also an inheritance effect. Thus, maybe what we observe here could be a superposition of two types of inheritance mechanisms and quantitatively unraveling the contribution of each will be a challenging task. Lastly, that salicylate induces inheritance effects in the activity of the *mar* operon should pique the interest of clinicians who assess the effects of aspirin on the *mar* operon induction in microbiome residents.

## Acknowledgements

Financial support came from the following sources: NIH P50 award P50GM081892-02 to the University of Chicago, a catalyst grant from the Chicago Biomedical Consortium with support from The Searle Funds at The Chicago Community Trust to PC, and a Yen Fellowship to CCG. We thank Raluca Vescan for her kind help with image processing software. We thank Wendy Grus and Kathleen Dave for editorial assistance.

## Author contributions

CCG and PC designed research; CCG, LB, PO performed research; CCG, LB, PO, MA, PC analyzed data; and CCG, LB and PC wrote the paper.

---

**Supplementary Figure S1.**
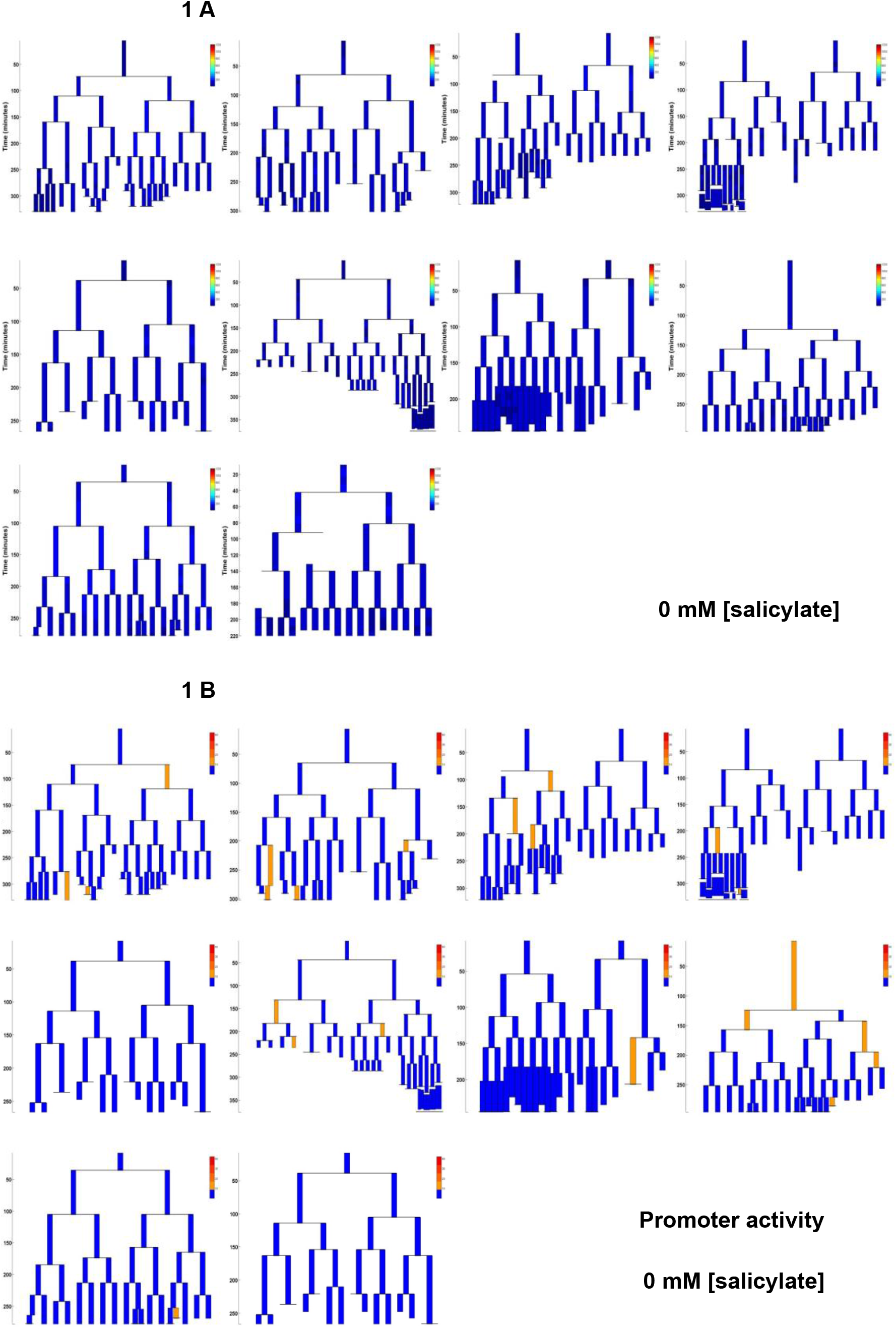
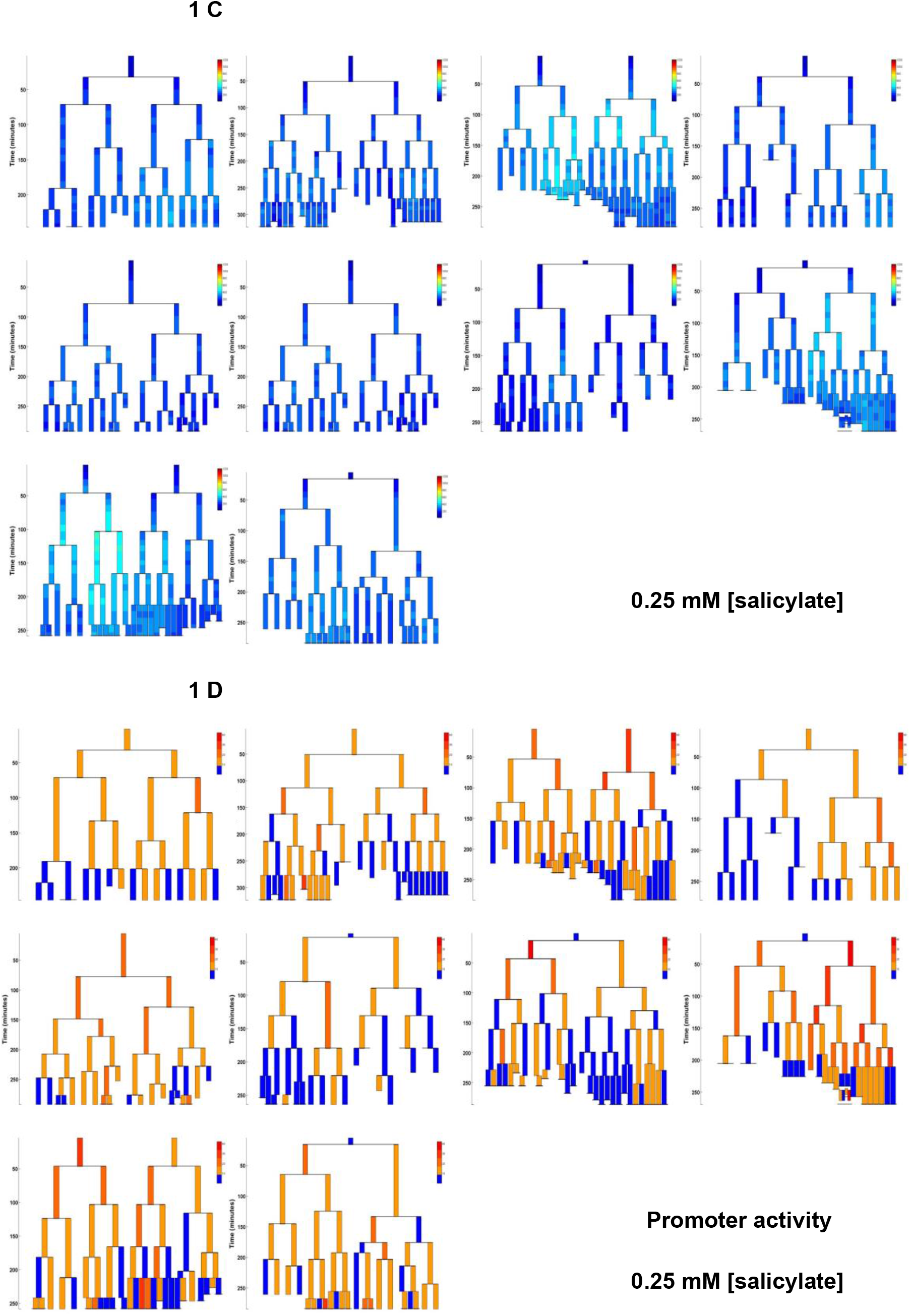
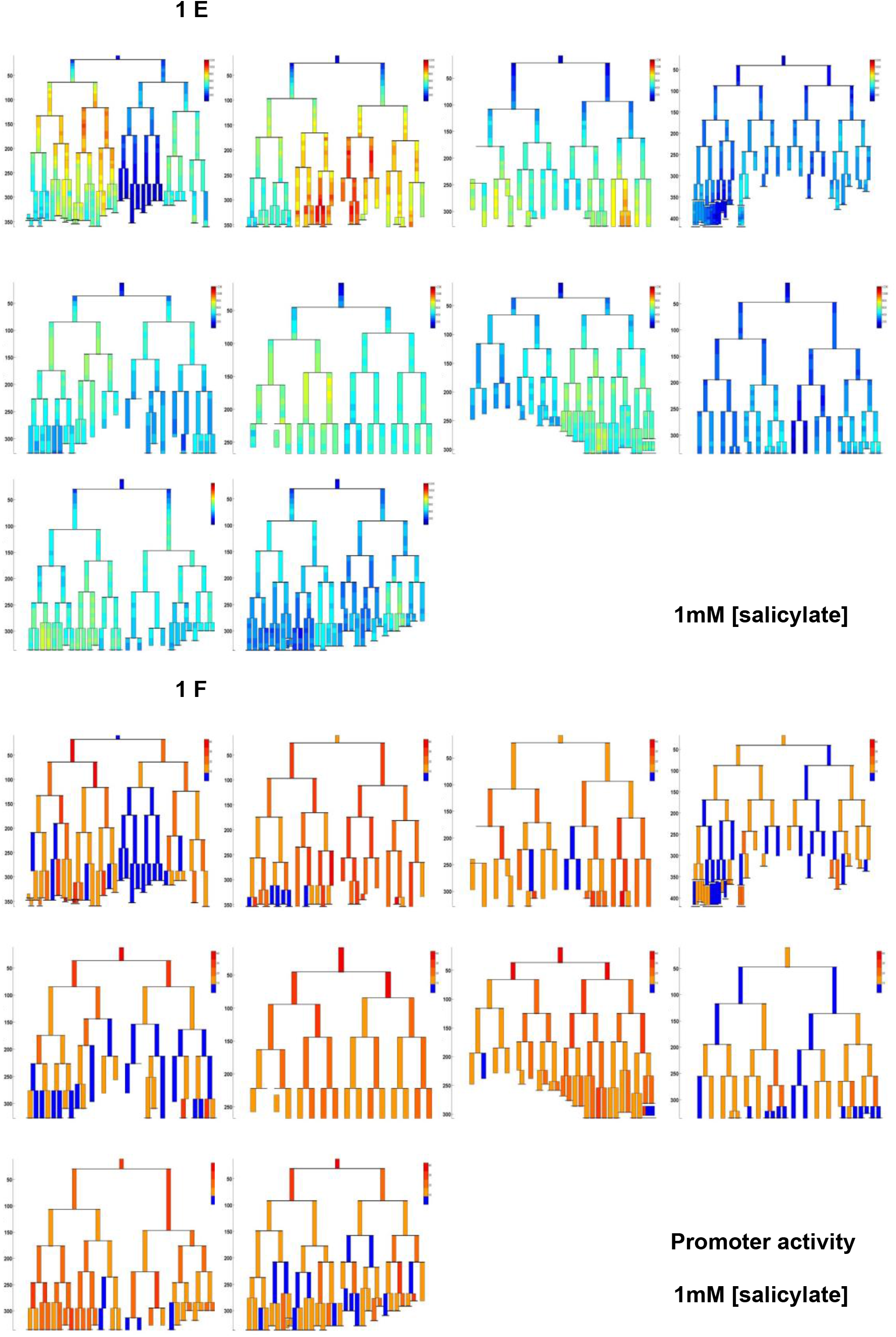
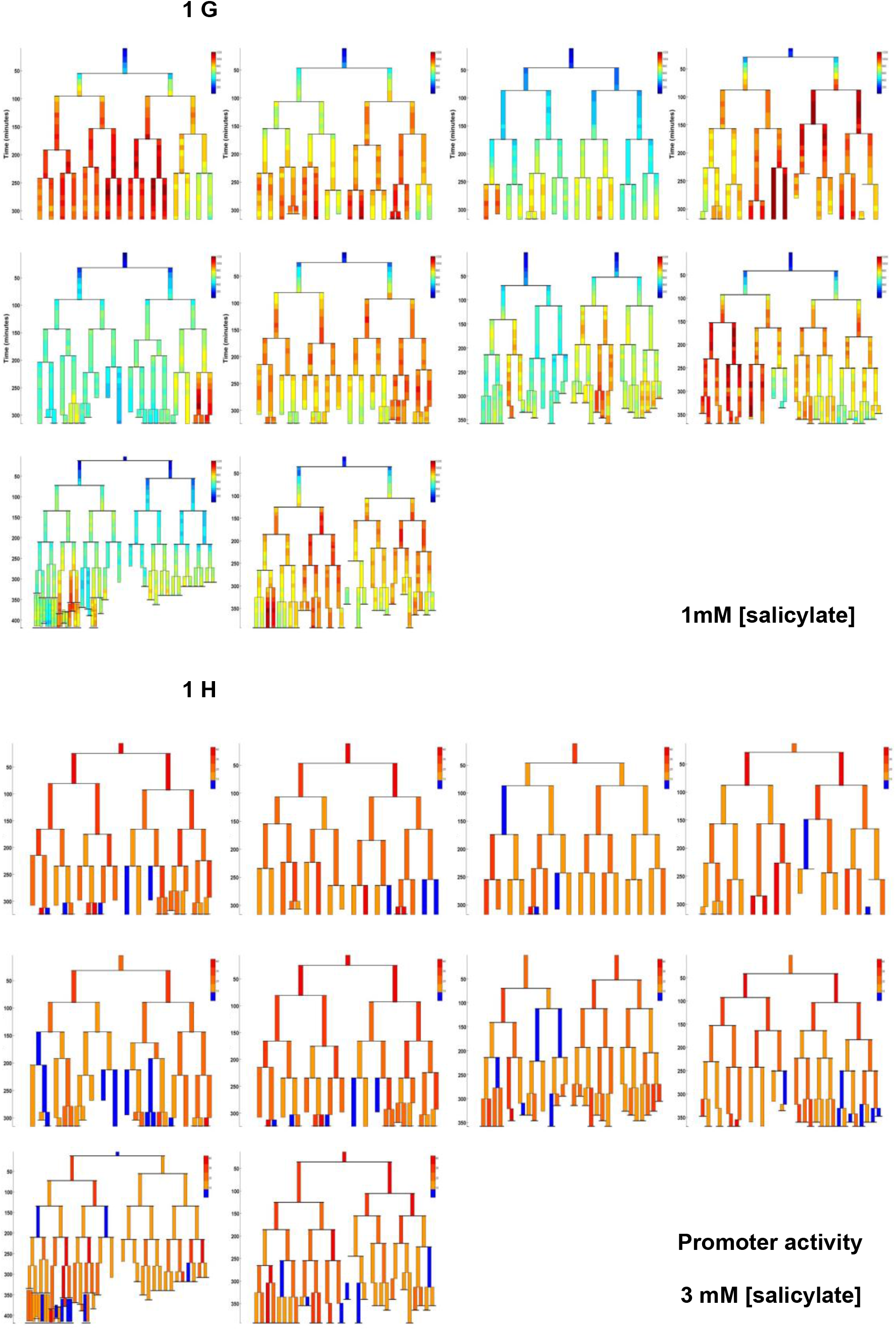
Complete sets of genealogy trees of *mar* promoter YFP levels and activities. **(a)** *mar* promoter YFP levels of genealogy trees for 0mM salicylate **(b)** Binarized *mar* promoter activity of genealogy trees for 0mM salicylate. These are the same trees as in (a) but with binarized values to display active and inactive cells. **(c)** *mar* promoter YFP levels of genealogy trees for 0.25mM salicylate **(d)** Binarized *mar* promoter activity of genealogy trees for 0mM salicylate. These are the same trees as in (c) but with binarized values to display active and inactive cells. **(e)** *mar* promoter YFP levels of genealogy trees for 1mM salicylate **(f)** Binarized *mar* promoter activity of genealogy trees for 0mM salicylate. These are the same trees as in (e) but with binarized values to display active and inactive cells. **(g)** *mar* promoter YFP levels of genealogy trees for 3mM salicylate **(h)** Binarized *mar* promoter activity of genealogy trees for 0mM salicylate. These are the same trees as in (g) but with binarized values to display active and inactive cells.

**Supplementary Figure S2.**
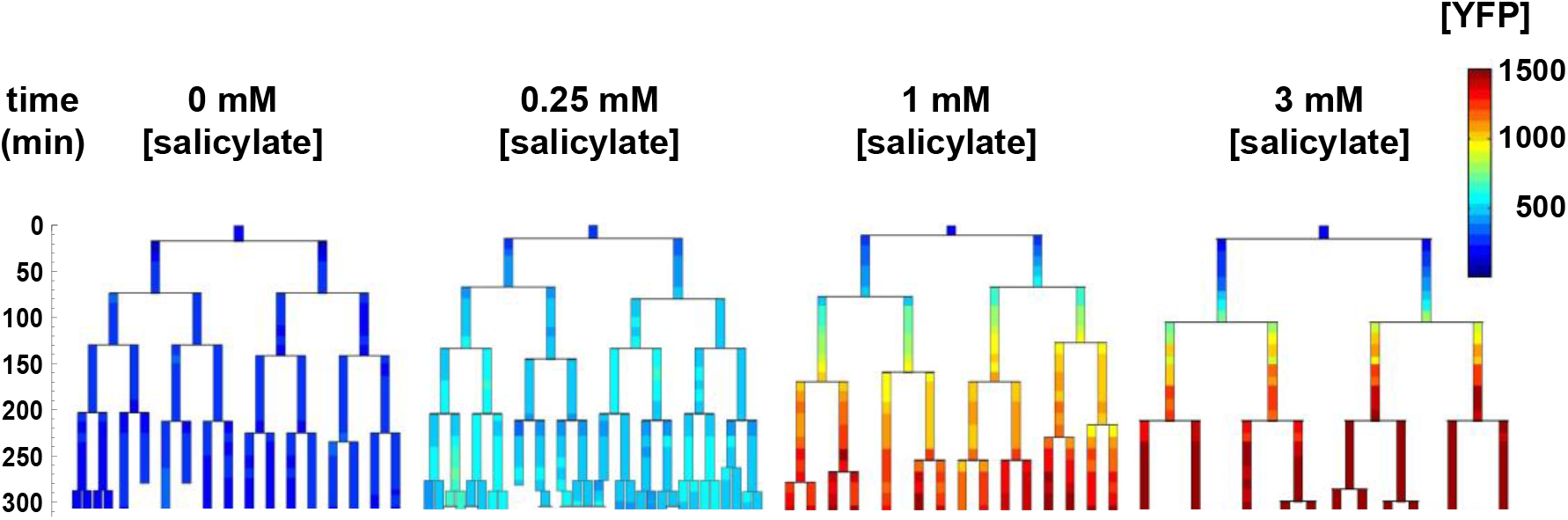
Genealogy trees of *mar* promoter induction in *ΔtolC* cells. Single cells with a plasmid carrying *yfp-venus* regulated by the *mar* promoter were monitored through several generations in the presence of the inducer salicylate. The genealogy tree follows the induced level of YFP concentration through the generations of a colony. Constant salicylate exposure began at time *t* = 0. Heat-map scale uses arbitrary fluorescence units to show low (blue) to high (red) [YFP].

## References

Alekshun MN, Levy SB. Regulation of chromosomally mediated multiple antibiotic resistance: The mar regulon. Antimicrobial Agents and Chemotherapy. 1997;41:2067–2075

Aracena J. Maximum number of fixed points in regulatory boolean networks. Bulletin of Mathematical Biology. 2008;70:1398–1409

Austin DW, Allen MS, McCollum JM, Dar RD, Wilgus JR, Sayler GS, Samatova NF, Cox CD, Simpson ML. Gene network shaping of inherent noise spectra. Nature. 2006;439:608–611

Baba T, Ara T, Hasegawa M, Takai Y, Okumura Y, Baba M, Datsenko KA, Tomita M, Wanner BL, Mori H. Construction of Escherichia coli K-12 in-frame, single-gene knockout mutants: The Keio collection. Molecular Systems Biology. 2006;2

Balaban NQ, Merrin J, Chait R, Kowalik L, Leibler S. Bacterial persistence as a phenotypic switch. Science. 2004;305:1622–1625

Balleza E, Kim JM, Cluzel P. Systemtic characterization of maturation time of fluorescent proteins in living cells. Nature Methods. 2018; 15: 47–51

Barbosa TM, Pomposiello PJ, The mar Regulon. Frontiers in Antimicrobial Resistance: A tribute to Stuart B. Levy 2005. Chapter 15

Bennett MR, Hasty J. Modelling microfluidic devices for measuring gene network dynamics in single cells. Nat Rev Genet. 2009;10:628–638

Bergmiller T, Andersson AMC, Tomasek K, Balleza E, Kiviet DJ, Hauschild R, Tkacik G, Guet CC. Biased partitioning of the multidrug efflux pump AcrAB-TolC underlies long-lived phenotypic heterogeneity. Science 2017; 356:311–315

Chang C, Garcia-Alcala M, Saiz L, Vilar JMG, Cluzel P. Robustness of DNA looping across multiple cell divisions in individual bacteria. PNAS 2022;119;e2200061119

Cluzel P, Surette M, Leibler S. An ultrasensitive bacterial motor revealed by monitoring signaling proteins in single cells. Science. 2000;287:1652–1655

Cohen SP, Levy SB, Foulds J, Rosner JL. Salicylate induction of antibiotic resistance in escherichia coli: Activation of the mar operon and a mar-independent pathway. J Bacteriol. 1993;175:7856–7862

Elowitz MB, Leibler S. A synthetic oscillatory network of transcriptional regulators. Nature. 2000;403:335–338

George AM, Levy SB. Amplifiable resistance to tetracycline, chloramphenicol, and other antibiotics in escherichia coli: Involvement of a non-plasmid-determined efflux of tetracycline. J Bacteriol. 1983;155:531–540

Guet CC, Elowitz MB, Hsing WH, Leibler S. Combinatorial synthesis of genetic networks. Science. 2002;296:1466–1470

Guet CC, Bruneaux L, Min TL, Siegal-Gaskins D, Figueroa I, Emonet T, Cluzel P. Minimally invasive determination of mrna concentration in single living bacteria. Nucleic Acids Research. 2008;36:-

Le TT, Harlepp S, Guet CC, Dittmar K, Emonet T, Pan T, Cluzel P. Real-time RNA profiling within a single bacterium. PNAS. 2005;102:9160–9164

Le TT, Emonet T, Harlepp S, Guet CC, Cluzel P. Dynamical determinants of drug-inducible gene expression in a single bacterium. Biophys J. 2006;90:3315–3321

Lewis K. Persister cells, dormancy and infectious disease. Nat Rev Microbiol. 2007;5:48–56

Li XZ, Nikaido H. Efflux-mediated drug resistance in bacteria. Drugs. 2004;64:159–204

Locke JCW, Elowitz MB. Using movies to analyse gene circuit dynamics in single cells. Nat Rev Microbiol. 2009;7:383–392

Lutz R, Bujard H. Independent and tight regulation of transcriptional units in escherichia coli via the lacR/o, the tetR/o and arac/i-1-i-2 regulatory elements. Nucleic Acids Research. 1997;25:1203–1210

Martin RG, Jair KW, Wolf RE, Rosner JL. Autoactivation of the marrab multiple antibiotic resistance operon by the mara transcriptional activator in escherichia coli. J Bacteriol. 1996;178:2216–2223

Martin RG, Rosner JL. Transcriptional and translational regulation of the MarRAB multiple antibiotic resistance operon in escherichia coli. Mol Microbiol. 2004;53:183–191

Martin RG, Bartlett ES, Rosner JL, Wall ME. Activation of the escherichia coli mara/soxs/rob regulon in response to transcriptional activator concentration. J Mol Biol. 2008;380:278–284

Nagai T, Ibata K, Park ES, Kubota M, Mikoshiba K, Miyawaki A. A variant of yellow fluorescent protein with fast and efficient maturation for cell-biological applications. Nature Biotechnology. 2002;20:87–90

Novick A, Weiner M. Enzyme induction as an all-or-none phenomenon. Proc Natl Acad Sci U S A. 1957;43:553–566

Rosner JL. Nonheritable resistance to chloramphenicol and other antibiotics induced by salicylates and other chemotactic repellents in Escherichia coli K-12. PNAS 1985;82:8771–8774

Siegele DA, Hu JC. Gene expression from plasmids containing the araBAD promoter at subsaturating inducer concentrations represents mixed populations. PNAS 1997; 94:8168–72

Stauffer D, Aharony A. Introduction to percolation theory. CRC Press; 1994

Taniguchi Y, Choi PJ, Li GW, Chen H, Babu M, Hearn J, Emili A, Xie XS. Quantifying e. Coli proteome and transcriptome with single-molecule sensitivity in single cells. Science. 2010;329:533–538

Thomas R, Richelle J. Positive feedback loops and multistationarity. Discrete Applied Mathematics. 1988;19:381–396

Vilar JMG, Guet CC, Leibler S. Modeling network dynamics: the lac operon, a case study. J Cell Bio 2003;161:471–76.

Zaslaver A, Bren A, Ronen M, Itzkovitz S, Kikoin I, Shavit S, Liebermeister W, Surette MG, Alon U. A comprehensive library of fluorescent transcriptional reporters for escherichia coli. Nat Methods. 2006;3:623–628

